# Epitope - based peptide vaccine against glycoprotein G of *Nipah henipavirus* using immunoinformatics approaches

**DOI:** 10.1101/678664

**Authors:** Arwa A. Mohammed, Shaza W. Shantier, Mujahed I. Mustafa, Hind K. Osman, Hashim E. Elmansi, Isam-Aldin A. Osman, Rawan A. Mohammed, Fatima A. Abdelrhman, Mihad E. Elnnewery, Einas M. Yousif, Marwa M.Mustafa, Nafisa M. Elfadol, Alaa I. Abdalla, Eiman Mahmoud, Ahmed A. Eltay, yassir A. Ahmed, Mohamed A. Hassan

## Abstract

**Background:** Nipah virus (NiV) is a member of the genus Henipavirus of the family Paramyxoviridae, characterized by high pathogenicity and endemic in South Asia, first emerged in Malaysia in 1998. The case-fatality varies from 40% to 70% depending on the severity of the disease and on the availability of adequate healthcare facilities. At present no antiviral drugs are available for NiV disease and the treatment is just supportive. Clinical presentation ranges from asymptomatic infection to fatal encephalitis. Bats are the main reservoir for this virus, which can cause disease in humans and animals. The last investigated NiV outbreak has occurred in May 2018 in Kerala.

**Objective:** This study aims to predict effective epitope-based vaccine against glycoprotein G of Nipah henipavirus using immunoinformatics approaches.

**Methods and Materials:** Glycoprotein G of Nipah henipavirus sequence was retrieved from NCBI. Different prediction tools were used to analyze the nominee’s epitopes in BepiPred-2.0: Sequential B-Cell Epitope Predictor for B-cell, T-cell MHC class II & I. Then the proposed peptides were docked using Autodock 4.0 software program.

**Results and Conclusions:** Peptide TVYHCSAVY shows a very strong binding affinity to MHC I alleles while FLIDRINWI shows a very strong binding affinity to MHC II and MHC I alleles. This indicates a strong potential to formulate a new vaccine, especially with the peptide FLIDRINWI that is likely to be the first proposed epitope-based vaccine against glycoprotein G of Nipah henipavirus. This study recommends an in-vivo assessment for the most promising peptides especially FLIDRINWI.

## 1. INTRODUCTION

Nipah virus (NiV) is an RNA virus that belongs to the Genus Henipavirus within the family Paramyxoviridae and has first emerged in Malaysia in 1998, gaining its name from a village called Sungai Nipah where it was isolated from the cerebrospinal fluid (CSF) of one of the patients ^[1–4]^. NiV is transmitted by zoonotic (from bats to humans, or from bats to pigs, and then to humans) as well as human-to-human routes. Its clinical presentation varies from asymptomatic (subclinical) infection to acute respiratory illness and fatal encephalitis with most of the patients have been in direct contact with infected pigs, it has also been found that the virus causes central nervous system illnesses in pigs and respiratory illnesses in horses resulting in a significant economic loss for farmer ^[1, 5–9]^. Large fruit bats of the genus Pteropus seem to act as a natural reservoir of NiV based on the isolation of Hendra virus which showed the presence of neutralizing antibodies to the Hendra virus on the bats ^[10, 11]^. Although there are no more cases of NiV in Malaysia, the outbreaks have been frequently occurring in India, Bangladesh, Thailand, and Cambodia ^[12]^. The case fatality rate ranges from 50 to 100%, making them one of the deadliest viruses known to infect human ^[3, 13, 14]^.

Laboratory diagnosis of Nipah virus infection is made using reverse transcriptase polymerase chain reaction (RT-PCR) from throat swabs, cerebrospinal fluid, urine, and blood analysis during acute and convalescent stages of the disease. IgG and IgM antibody detection can be done after recovery to confirm Nipah virus infection. Immunohistochemistry on tissues collected during autopsy also confirms the disease ^[15, 16]^. Currently, there is no effective treatment for the Nipah Virus infection however a few precautions include practicing standard infection control, barrier nursing to avoid the spread of infection from person to person as well as the isolation of those suspected to have the infection ^[7, 8, 17]^. Recent computational approaches have provided further information about viruses, including the study conducted by Badawi M, et al. on ZIKA virus, where the envelope glycoprotein was obtained using protein databases. The most immunogenic epitope for the T and B cells involved in cell-mediated immunity were previously analyzed ^[18]^. The main focus of the analysis was the MHC class-I potential peptides using in silico analysis techniques ^[19, 20]^. In this study, the same techniques were applied to keep MHC class I and II along with the world population coverage as our main focus. Furthermore, we aim to design an Epitope-Based Peptide Vaccine against Nipah virus using peptides of its glycoprotein G as an immunogenic part to stimulate a protective immune response ^[3]^.

Nipah virus invades host cells by the fusion of the host cell membranes at physiological pH without requiring viral endocytosis. Cell-cell fusion is a pathological lineament of Nipah virus infections, resulting in cell-to-cell spread, inflammation, and destruction of endothelial cells and neurons ^[21]^. Both Nipah virus entry and Cell-cell fusion require the concerted efforts of the attachment of glycoprotein G and fusion (F) glycoprotein. Upon receptor binding, Nipah virus glycoprotein G triggers a conformational cascade in Nipah virus glycoprotein F that executes viral and/or cell membrane fusion ^[22]^. Due, to the potency of glycoprotein G over F, we have considered it as the target of this study. It could be the first vaccine developed for humans against glycoprotein G of *Nipah henipavirus* to be put forward using an immunoinformatics approach and population coverage analytical tools. Even though there’s a lot of challenges regarding the development of peptide vaccines, we have decided to develop them for fighting the Nipah virus infection because they make a very good alternative strategy that relies on the usage of short peptide fragments to induce immune responses that are extremely targeted, avoiding all allergenic as well as reactogenic sequences ^[23–26]^. Antigenic epitopes from single proteins may not be really necessary, whereas some of these epitopes may even be detrimental to the induction of protective immunity. This logic has created an interest in peptide vaccines and especially those containing only epitopes that are capable of inducing desirable T cell and B cell mediated immune response. Less than 20 amino acid sequences make up the peptides used in such vaccines, which are then synthesized to form an immunogenic peptide molecule. These molecules represent the specific epitope of an antigen. These vaccines are also capable of inducing immunity against different strains of a specific pathogen by forming non-contiguous and immunodominant epitopes that are usually conserved in the strains of the pathogen ^[27]^.

The production of peptide vaccines is extremely safe and cost-effective especially when they are compared to conventional vaccines. Traditional vaccines that prevent emerging infectious diseases (EIDs) are very difficult to produce because they require the need to culture pathogenic viruses in vitro however, epitope based peptide vaccines do not require any means of in vitro culturing making them biologically safe, they allow for the large scale of bio-processing to be carried out rapidly and economically and finally their selectivity allows for the precise activation of immune responses by means of selecting immunodominant and conserved epitopes ^[25, 28]^. The complexity of the epitope based peptide vaccine’s design depends largely on the properties of the carrier molecules from their reactogenicity to their allergenicity ^[29, 30]^. When it comes to the selection of epitopes, it is directed towards B cells, cytotoxic T cells and the induction of the helper T cells as well. Then, it is important to identify the epitopes capable of activating T cells vital for bringing about protective immunity. One of the issues concerned with peptide vaccines representing T cells in a human population that is highly MHC- heterogeneous is identifying highly conserved immunodominant epitopes which are considered to be broad-spectrum vaccines due to their ability to work against multiple serovars of a pathogen ^[30]^.

For the purpose of this research, we have used a variety of bioinformatics tools for the prediction of epitopes along with the population coverage and epitope selection algorithms including the translocation of peptides into MHC class I and MHC class II.

## 2. Materials and Methods

**Figure 1:**
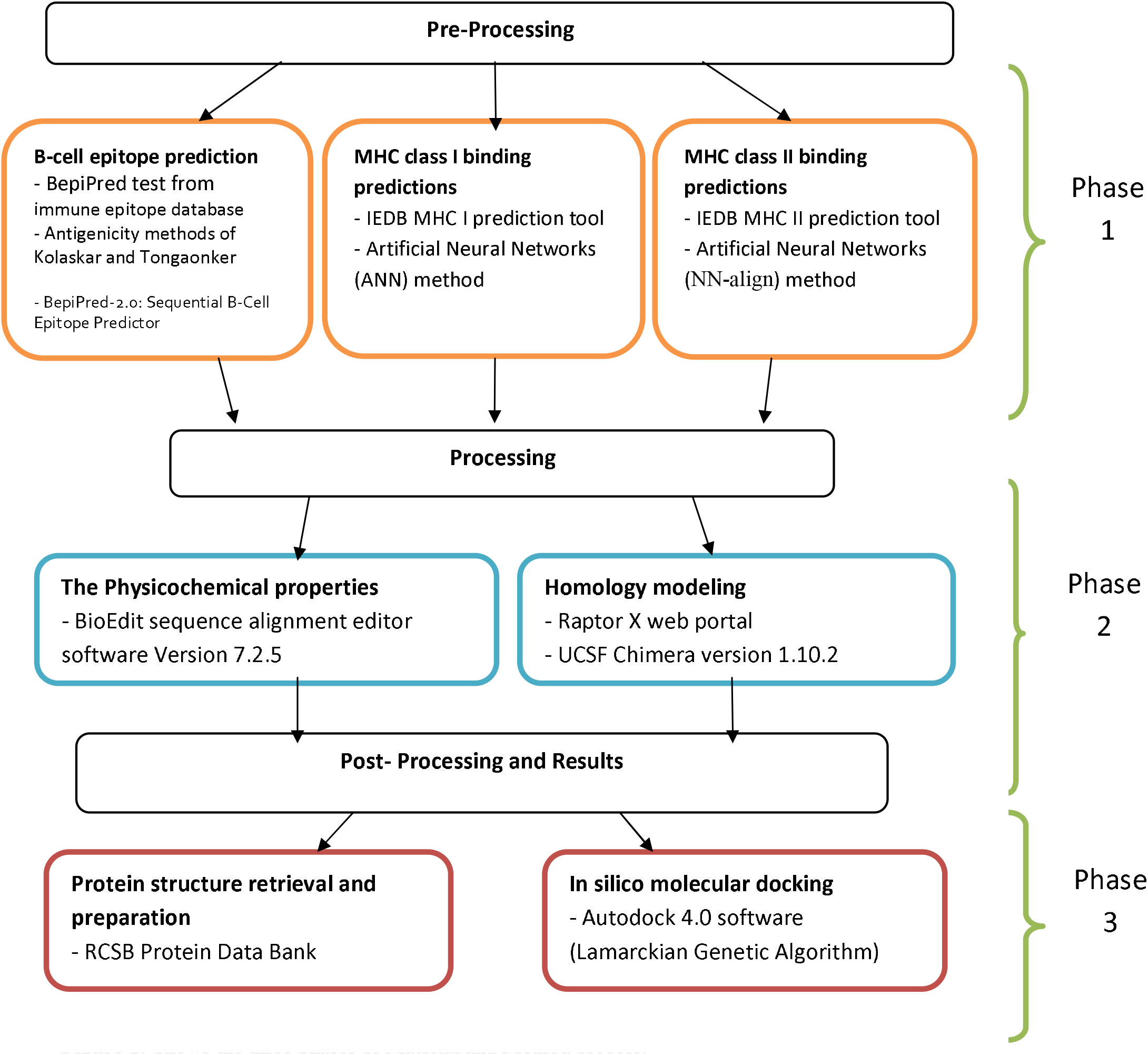
Shows the three phases of Material and Method process.

### 2.1 Sequences retrieval

The amino acids sequences of Glycoprotein G (Glycoside hydrolase family) for a total of 21 strains of Nipah virus were retrieved from NCBI database (https://www.ncbi.nlm.nih.gov/protein) ^[31]^ in FASTA format on July 2018. Different prediction tools of Immune Epitope Database IEDB analysis resource (http://www.iedb.org/) ^[32]^ were then used to analyze the candidate epitopes.

### 2.2 Conservation region and physicochemical properties

Conservation regions were determined using multiple sequence alignment with the help of Clustal-W in the Bio-edit software version ^[33]^. Epitope conservancy prediction for individual epitopes was then calculated using the IEDB analysis resource. Conservancy can be defined as the portion of protein sequences that restrain the epi-tope measured at or exceeding a specific level of identity ^[34]^. The physicochemical properties of the retrieved sequence; molecular weight and amino acid composition; were also determined by Bio-edit software version.

### 2.3 B cell epitope prediction tools

Candidate epitopes were analyzed using several B-cell prediction methods to determine the antigenicity, flexibility, hydrophilicity and surface accessibility. The linear prediction epitopes were obtained from Immune epitope database (http://tools.iedb.org/bcell/result/) ^[35]^ by using BepiPred test with a threshold value of 0.149 and a window size 6.0

Moreover, surface accessible epitopes were predicated with a threshold value of 1.0 and window size 6.0 using the Emini surface accessibility prediction tool ^[35]^.

Kolaskar and Tongaonker antigenicity methods (http://tools.iedb.org/bcell/result/) were proposed to determine the sites of antigenic epitopes with a default threshold value of 1.030 and a window size 6.0 ^[36]^.

### 2.4 T cell epitope prediction tools

#### 2.4.1 Peptide binding to MHC class 1 molecules

The peptide binding was assessed by the IEDB MHC 1 prediction tool at http://tools.iedb.org/mhc1. This tool employs different methods to determine the ability of the submitted sequence to bind to a specific MHC class 1 molecule. The artificial neural network (ANN) method was used to calculate IC50 values of peptide binding to MHC-1 molecules. For both frequent and non-frequent alleles, peptide length was set to 9 amino acids earlier to the prediction. The alleles having binding affinity IC50 equal to or less than 500 nM were considered for further analysis ^[37]^.

#### 2.4.2 Peptide binding to MHC class II molecules

To predict the peptide binding to MHC class II molecules, MHC II prediction tool http://tools.iedb.org/mhcII provided by Immune Epitope Database (IEDB) analysis resource and human allele references set was used ^[38]^. The Artificial Neural Network prediction method was chosen to identify the binding affinity to MHC II grooves and MHC II binding core epitopes. All epitopes that bind to many alleles at score equal to or less than 1000 half-maximal inhibitory concentration (IC50) were selected for further analysis.

### 2.5 Population coverage

The population coverage of each epitope was calculated by IEDB population coverage tool at (http://tools.iedb.org/tools/population/iedb_input) This tool is aimed in order to determine the fraction of individuals predicted to respond to a given set of epitopes with known MHC restrictions ^[39]^. For every single population coverage, the tool computed the following information: (1) predicted population coverage, (2) HLA combinations recognized by the population, and (3) HLA combinations recognized by 90% of the population (PC90). All epitopes and their MHCI and MHC II molecules were assessed against population coverage area selected before submission.

### 2.6 Homology modeling

The 3D structure of glycoprotein G of Nipah virus was predicted using the Raptor X web portal (http://raptorx.uchicago.edu/) the reference sequence was submitted in FASTA format on 14/9/2018 and structure received on 15/9/2018 ^[40]^. The Structure was then treated with UCSF Chimera 1.10.2 to visualize the position of proposed peptides ^[41]^.

### 2.7 In silico Molecular Docking

#### 2.7.1 Ligand Preparation

In order to estimate the binding affinities between the epitopes and molecular structure of MHC I & MHC II, in silico molecular docking were used. Sequences of proposed epitopes were selected from Nipah virus reference sequence using Chimera 1.10 and saved as a pdb file. The obtained files were then optimized and energy minimized. The HLA-A0201 was selected as the macromolecule for docking. Its crystal structure (4UQ3) was downloaded from the RCSB Protein Data Bank (http://www.rcsb.org/pdb/home/home.do), which was in a complex with an azobenzene-containing peptide ^[42]^.

All water molecules and heteroatoms in the retrieved target file 4UQ3 were then removed. Target structure was further optimized and energy minimized using Swiss PDB viewer V.4.1.0 software ^[43]^.

Molecular docking was performed using Autodock 4.0 software, based on Lamarckian Genetic Algorithm; which combines energy evaluation through grids of affinity potential to find the suitable binding position for a ligand on a given protein ^[44,45]^ Polar hydrogen atoms were added to the protein targets and Kollman united atomic charges were computed. The target’s grid map was calculated and set to 60×60×60 points with grid spacing of 0.375 Ǻ. The grid box was then allocated properly in the target to include the active residue in the center. The genetic algorithm and its run were set to 100. The docking algorithms were set to default. Finally, results were retrieved as binding energies and poses that showed lowest binding energies were visualized using UCSF chimera.

## 3. Results

### 3.1. *Nipah virus glycoprotein G* physical and chemical parameters

The physicochemical properties of *Nipah virus glycoprotein G* protein were assessed using BioEdit software version 7.0.9.0. The protein length was found to be 602 amino acids and the molecular weight was 67035.54 Daltons. The amino acid that formed *Nipah virus glycoprotein G* protein and their number along with their molar percentage (Mol%) were shown in table (1).

**Figure 2:**
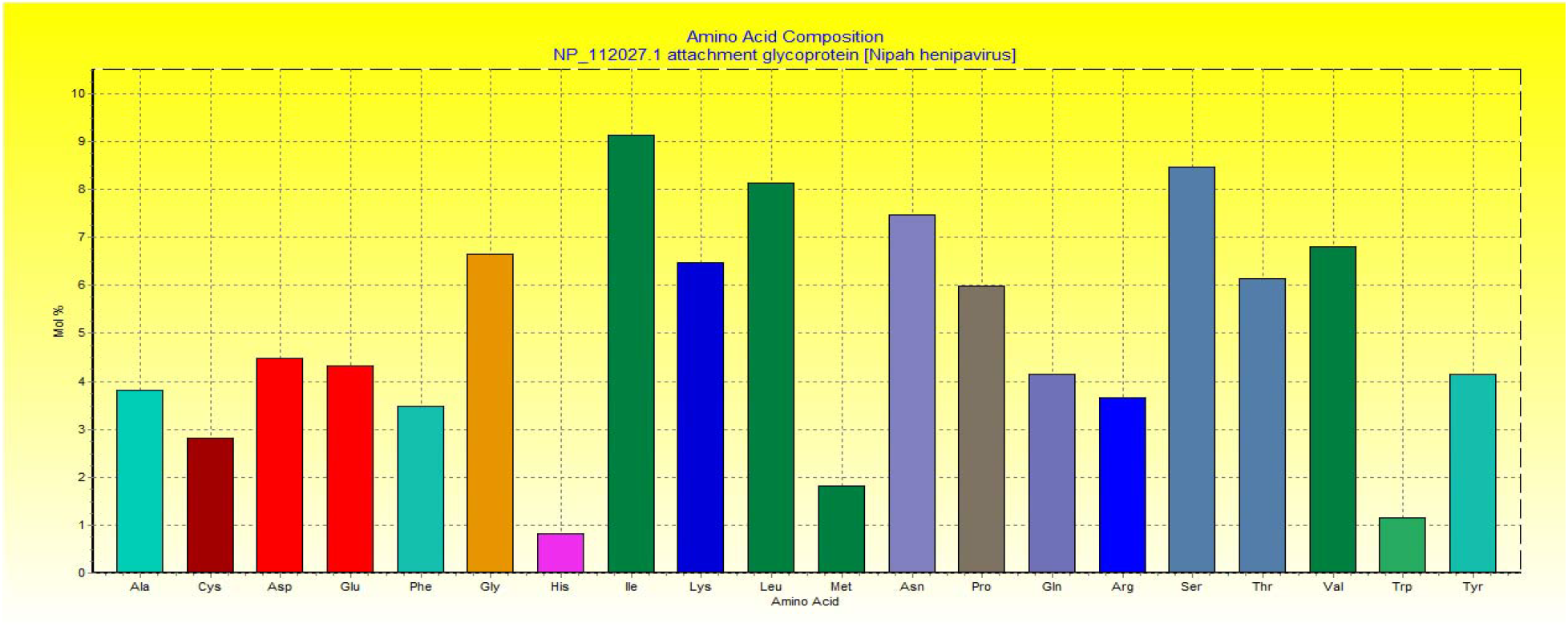
Amino acids composition of Nipah henipavirus glycoprotein G using Bio Edit software Version 7.2.5.

**Table 1:** Number and Mol% of amino acids constituted *Nipah virus glycoprotein G* using Bio Edit software Version 7.2.5.

| Amino Acid | Number | Mol% | Amino Acid | Number | Mol% |
| --- | --- | --- | --- | --- | --- |
| Ala A | 23 | 3.82 | Met M | 11 | 1.83 |
| Cys C | 17 | 2.82 | Asn N | 45 | 7.48 |
| Asp D | 27 | 4.49 | Pro P | 36 | 5.98 |
| Glu E | 26 | 4.32 | Gln Q | 25 | 4.15 |
| Phe F | 21 | 3.49 | Arg R | 22 | 3.65 |
| Gly G | 40 | 6.64 | Ser S | 51 | 8.47 |
| His H | 5 | 0.83 | Thr T | 37 | 6.15 |
| Ile I | 55 | 9.14 | Val V | 41 | 6.81 |
| Lys K | 39 | 6.48 | Trp W | 7 | 1.16 |
| Leu L | 49 | 8.14 | Tyr Y | 25 | 4.15 |

### 3.2. B-cell epitope prediction

The sequence of *Nipah virus glycoprotein G* was subjected to Bepipred linear epitope prediction, Emini surface accessibility, and Kolaskar and Tongaonkar antigenicity methods in IEDB, to determine the binding to B cell, being in the surface and to test the immunogenicity. The results were shown in Figure (3-5).

**Figure 3:**
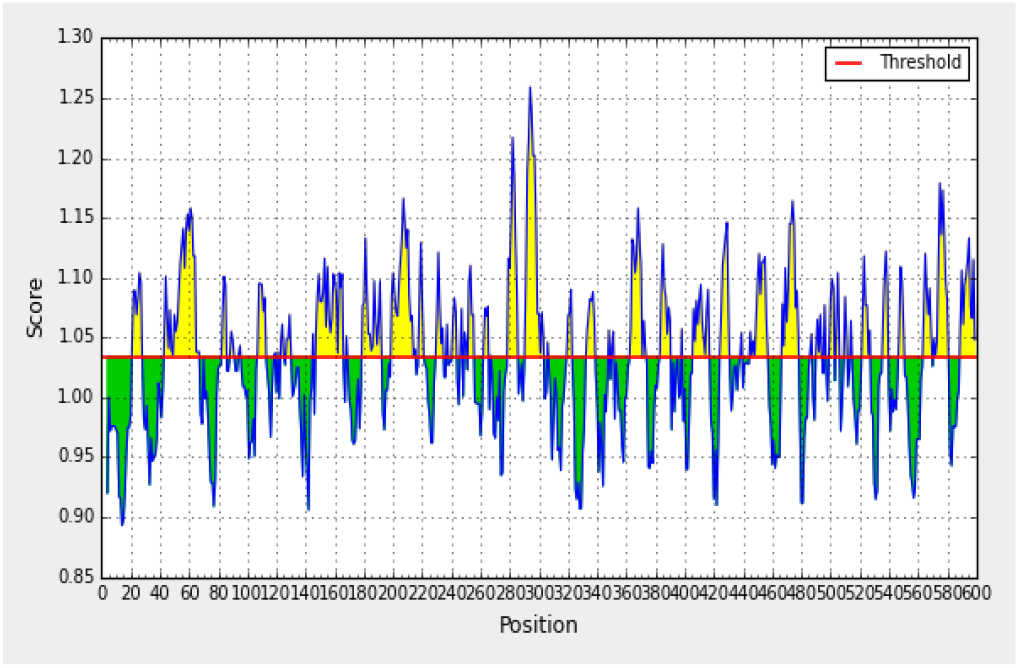
Represents Bepipred Linear Prediction; Areas above red line (threshold) are suggested to B cell epitope while green areas are not.

**Figure 4:**
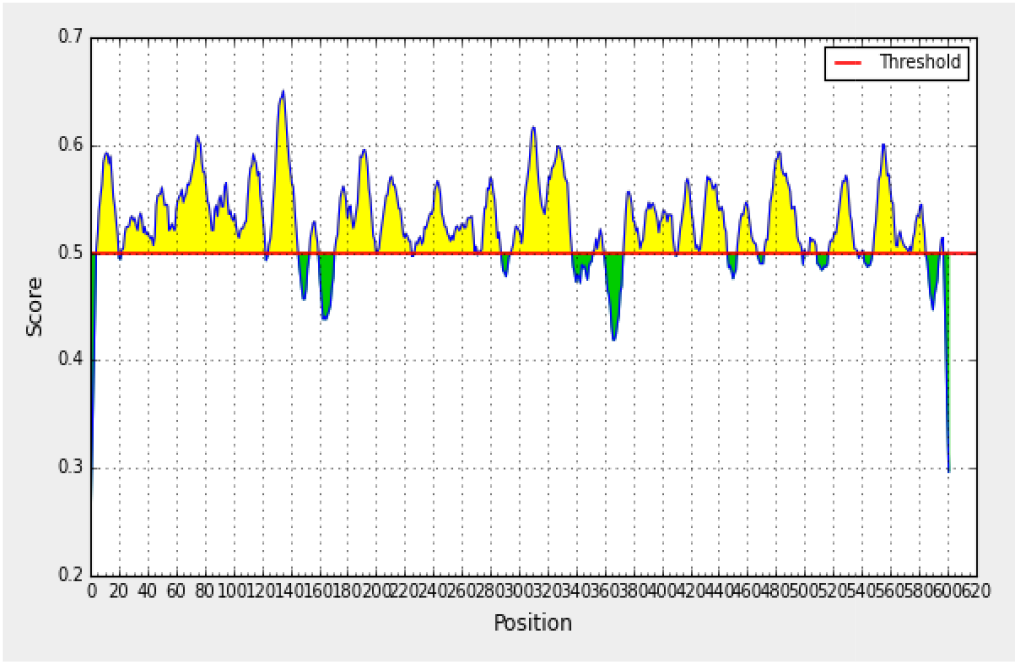
Emini surface accessibility prediction; Areas above red line (threshold) are suggested to B cell epitope while green areas are not.

**Figure 5:**
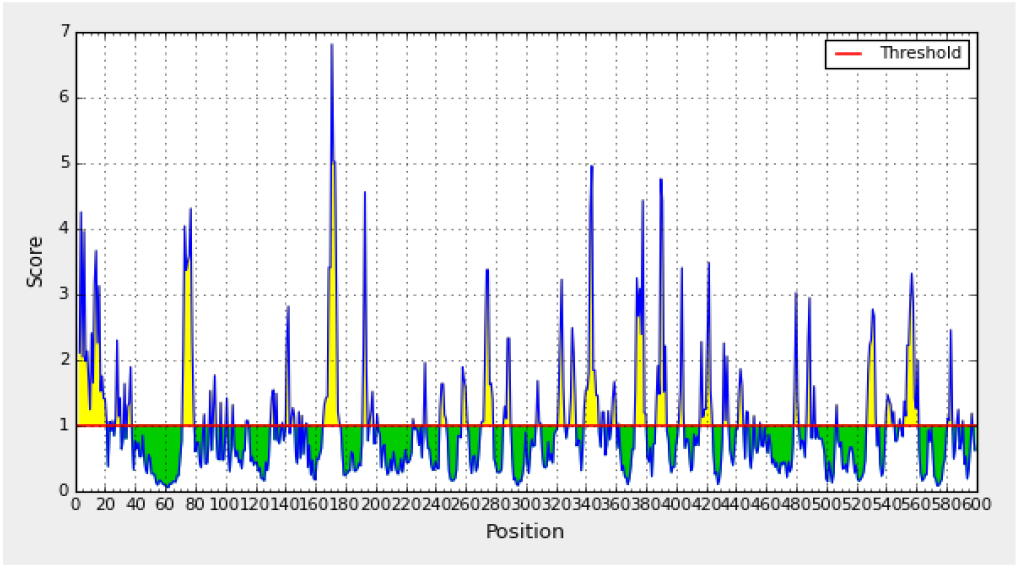
Kolaskar and Tongaonkar antigenicity prediction; Areas above red line (threshold) are suggested to B cell epitope while green areas are not.

### 3.3. Prediction of T helper cell epitopes and interaction with MHC I alleles

*Nipah virus glycoprotein G* sequence was analyzed using IEDB MHC I binding prediction tool based on ANN-align with half-maximal inhibitory concentration (IC_50_) ≤500; the list most promising epitopes that had Binding affinity with the Class I alleles along with their positions in the *Nipah virus glycoprotein G* were shown in table (2).

**Table 2.**
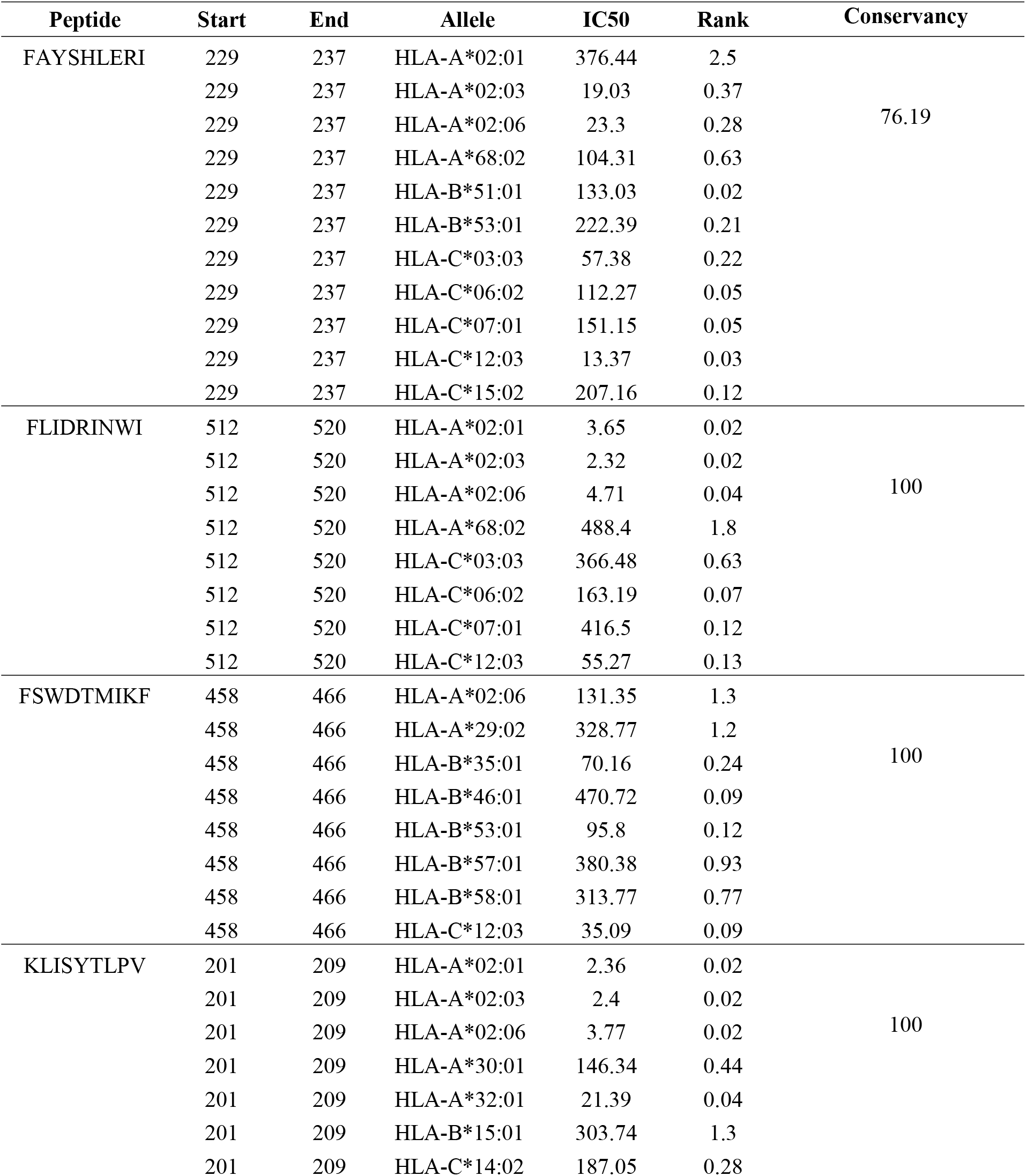
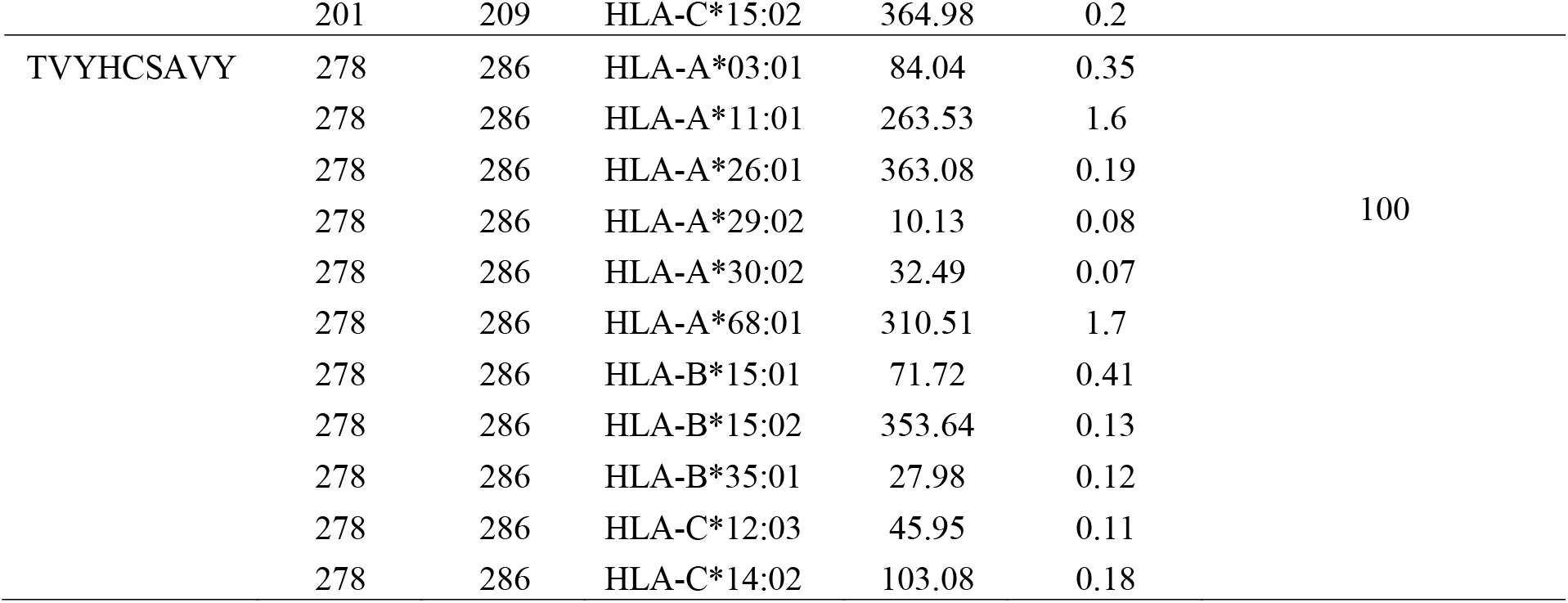
Most potential T-cell epitopes with interacting MHC-1 alleles, their positions, IC50, rank, and conservancy

### 3.4. Prediction of T helper cell epitopes and interaction with MHC II alleles

*Nipah virus glycoprotein G* sequence was analyzed using IEDB MHC- II binding prediction tool based on NN-align with half-maximal inhibitory concentration (IC_50_) ≤1000; the list of the most promising epitopes and their correspondent binding MHC11 alleles along with their positions in the *Nipah virus glycoprotein G* were shown in (supplementary table 1) while the list most promising epitopes that had a strongly Binding affinity with the Class II alleles and the number of their binding alleles were shown in table (3).

**Table 3:** Most potential T-cell epitopes (core sequence) that had a binding affinity with MHC II alleles and the number of their binding alleles.

| Core sequence | Alleles | Number of allele |
| --- | --- | --- |
| FLIDRINWI | HLA-DPA1*01/DPB1*04:01 | 12 |
|  | HLA-DPA1*01:03/DPB1*02:01 |  |
|  | HLA-DPA1*02:01/DPB1*01:01 |  |
|  | HLA-DPA1*02:01/DPB1*05:01 |  |
|  | HLA-DPA1*03:01/DPB1*04:02 |  |
|  | HLA-DQA1*01:01/DQB1*05:01 |  |
|  | HLA-DQA1*05:01/DQB1*02:01 |  |
|  | HLA-DRB1*01:01 |  |
|  | HLA-DRB1*03:01 |  |
|  | HLA-DRB1*04:01 |  |
|  | HLA-DRB1*04:04 |  |
|  | HLA-DRB1*04:05 |  |
| FAYSHLERI | HLA-DPA1*01:03/DPB1*02:01 | 13 |
|  | HLA-DPA1*02:01/DPB1*01:01 |  |
|  | HLA-DPA1*02:01/DPB1*05:01 |  |
|  | HLA-DPA1*03:01/DPB1*04:02 |  |
|  | HLA-DQA1*05:01/DQB1*02:01 |  |
|  | HLA-DQA1*05:01/DQB1*03:01 |  |
|  | HLA-DRB1*01:01<br>HLA-DRB1*04:04<br>HLA-DRB1*04:05<br>HLA-DRB1*07:01<br>HLA-DRB1*09:01<br>HLA-DRB3*01:01<br>HLA-DRB5*01:01 |  |
| FIEISDQRL | HLA-DPA1*01:03/DPB1*02:01<br>HLA-DPA1*02:01/DPB1*01:01<br>HLA-DPA1*03:01/DPB1*04:02<br>HLA-DQA1*05:01/DQB1*02:01<br>HLA-DRB1*01:01<br>HLA-DRB1*04:05<br>HLA-DRB1*07:01<br>HLA-DRB1*08:02<br>HLA-DRB1*13:02<br>HLA-DRB1*15:01<br>HLA-DRB4*01:01<br>HLA-DRB5*01:01<br>HLA-DRB1*04:01<br>HLA-DRB1*07:01<br>HLA-DRB1*09:01<br>HLA-DRB1*11:01<br>HLA-DRB1*13:02 | 17 |
| ILSAFNTVI | HLA-DPA1*03:01/DPB1*04:02<br>HLA-DQA1*05:01/DQB1*03:01<br>HLA-DRB1*01:01<br>HLA-DRB1*04:01<br>HLA-DRB1*04:05<br>HLA-DRB1*07:01<br>HLA-DRB1*08:02<br>HLA-DRB1*09:01<br>HLA-DRB1*11:01<br>HLA-DRB1*13:02<br>HLA-DRB1*15:01<br>HLA-DRB4*01:01<br>HLA-DRB5*01:01 | 13 |
| TVYHCSAVY | HLA-DQA1*05:01/DQB1*03:01<br>HLA-DRB1*07:01<br>HLA-DRB1*13:02<br>HLA-DRB1*15:01 | 4 |

### 3.5 Population coverage

Population coverage test was performed to detect all epitopes binds to MHC1 alleles and MHC11 alleles for the world, South Asia, Southeast Asia, Sudan, and North Africa.

**Table 4.** Population coverage average for all epitopes binding to MHC I alleles in World.

| Population/area | Class I |  |  |
| --- | --- | --- | --- |
|  | coverage<sup>a</sup> | average_hit<sup>b</sup> | pc90<sup>c</sup> |
| <u>World</u> | 99.84% | 36.62 | 16.87 |
| **Average** | **99.84** | **36.62** | **16.87** |
| **Standard deviation** | **0.0** | **0.0** | **0.0** |

<sup>a</sup> projected population coverage
<sup>b</sup> average number of epitope hits / HLA combinations recognized by the population
<sup>c</sup> minimum number of epitope hits / HLA combinations recognized by 90% of the population

**Table 5.** Population coverage average for all epitopes binding to MHC I alleles in South Asia

| Population/area | Class I |  |  |
| --- | --- | --- | --- |
|  | coverage<sup>a</sup> | average_hit<sup>b</sup> | pc90<sup>c</sup> |
| <u>South Asia</u> | 98.4% | 30.6 | 9.29 |
| **Average** | **98.4** | **30.6** | **9.29** |
| **Standard deviation** | **0.0** | **0.0** | **0.0** |

<sup>a</sup> projected population coverage
<sup>b</sup> average number of epitope hits / HLA combinations recognized by the population
<sup>c</sup> minimum number of epitope hits / HLA combinations recognized by 90% of the population

**Table 6.** Population coverage average for all epitopes binding to MHC I alleles in Southeast Asia

| Population/area | Class I |  |  |
| --- | --- | --- | --- |
|  | coverage<sup>a</sup> | average_hit<sup>b</sup> | pc90<sup>c</sup> |
| <u>Southeast Asia</u> | 98.55% | 28.42 | 8.61 |
| **Average** | **98.55** | **28.42** | **8.61** |
| **Standard deviation** | **0.0** | **0.0** | **0.0** |

<sup>a</sup> projected population coverage
<sup>b</sup> average number of epitope hits / HLA combinations recognized by the population
<sup>c</sup> minimum number of epitope hits / HLA combinations recognized by 90% of the population

**Table 7.** Population coverage average for all epitopes binding to MHC I alleles in Sudan

| Population/area | Class I |  |  |
| --- | --- | --- | --- |
|  | coverage<sup>a</sup> | average_hit<sup>b</sup> | pc90<sup>c</sup> |
| <u>Sudan</u> | 99.36% | 34.02 | 13.4 |
| **Average** | **99.36** | **34.02** | **13.4** |
| **Standard deviation** | **0.0** | **0.0** | **0.0** |

<sup>a</sup> projected population coverage<sup>b</sup> average number of epitope hits / HLA combinations recognized by the population<sup>c</sup> minimum number of epitope hits / HLA combinations recognized by 90% of the population**Table 8.** Population coverage average for all epitopes binding to MHC I alleles in North Africa

**Table 8.**
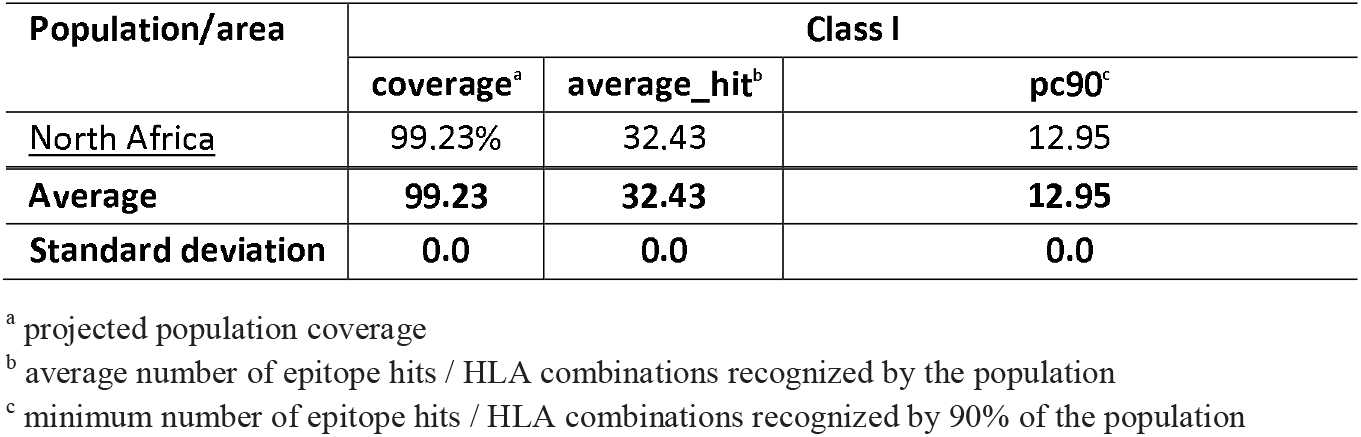
Population coverage average for all epitopes binding to MHC I alleles in North Africa

**Table 9:** Population coverage for all epitopes binding to MHC II in World

| Population/area | Class II |  |  |
| --- | --- | --- | --- |
|  | coverage<sup>a</sup> | average_hit<sup>b</sup> | pc90<sup>c</sup> |
| <u>World</u> | 56.84% | 54.88 | -24.24 |
| **Average** | **56.84** | **54.88** | **-24.24** |
| **Standard deviation** | **0.0** | **0.0** | **0.0** |

<sup>a</sup> projected population coverage<sup>b</sup> average number of epitope hits / HLA combinations recognized by the population<sup>c</sup> minimum number of epitope hits / HLA combinations recognized by 90% of the population**Table 10:** Population coverage for all epitopes binding to MHC II in Southeast Asia

**Table 10:** Population coverage for all epitopes binding to MHC II in Southeast Asia

| Population/area | Class II |  |  |
| --- | --- | --- | --- |
|  | coverage<sup>a</sup> | average_hit<sup>b</sup> | pc90<sup>c</sup> |
| <u>Southeast Asia</u> | 48.63% | 36.27 | 1.65 |
| **Average** | **48.63** | **36.27** | **1.65** |
| **Standard deviation** | **0.0** | **0.0** | **0.0** |

<sup>a</sup> projected population coverage<sup>b</sup> average number of epitope hits / HLA combinations recognized by the population<sup>c</sup> minimum number of epitope hits / HLA combinations recognized by 90% of the population

**Table 11:** Population coverage for all epitopes binding to MHC II in South Asia

| Population/area | Class II |  |  |
| --- | --- | --- | --- |
|  | coverage<sup>a</sup> | average_hit<sup>b</sup> | pc90<sup>c</sup> |
| <u>South Asia</u> | 56.0% | 50.5 | -10.09 |
| **Average** | **56.0** | **50.5** | **-10.09** |
| **Standard deviation** | **0.0** | **0.0** | **0.0** |

<sup>a</sup> projected population coverage<sup>b</sup> average number of epitope hits / HLA combinations recognized by the population<sup>c</sup> minimum number of epitope hits / HLA combinations recognized by 90% of the population**Table 12:** Population coverage for all epitopes binding to MHC II in Sudan

**Table 12:** Population coverage for all epitopes binding to MHC II in Sudan

| Population/area | Class II |  |  |
| --- | --- | --- | --- |
|  | coverage<sup>a</sup> | average_hit<sup>b</sup> | pc90<sup>c</sup> |
| <u>Sudan</u> | 55.75% | 36.89 | 4.6 |
| **Average** | **55.75** | **36.89** | **4.6** |
| **Standard deviation** | **0.0** | **0.0** | **0.0** |

<sup>a</sup> projected population coverage<sup>b</sup> average number of epitope hits / HLA combinations recognized by the population<sup>c</sup> minimum number of epitope hits / HLA combinations recognized by 90% of the population**Table 13:** Population coverage for all epitopes binding to MHC II in North Africa

**Table 13:** Population coverage for all epitopes binding to MHC II in North Africa

| Population/area | Class II |  |  |
| --- | --- | --- | --- |
|  | coverage<sup>a</sup> | average_hit<sup>b</sup> | pc90<sup>c</sup> |
| <u>North Africa</u> | 62.37% | 50.09 | -3.31 |
| **Average** | **62.37** | **50.09** | **-3.31** |
| **Standard deviation** | **0.0** | **0.0** | **0.0** |

<sup>a</sup> projected population coverage<sup>b</sup> average number of epitope hits / HLA combinations recognized by the population<sup>c</sup> minimum number of epitope hits / HLA combinations recognized by 90% of the population

**Table 14:** Population coverage of the three highly proposed peptides in MHCI, MHCII and both MHC I & II in five areas.

| Peptide | Population coverage %/ Area |  |  |  |  |  |  |  |  |  |  |  |  |  |  |
| --- | --- | --- | --- | --- | --- | --- | --- | --- | --- | --- | --- | --- | --- | --- | --- |
|  | World |  |  | Southeast Asia |  |  | South Asia |  |  | North Africa |  |  | Sudan |  |  |
|  | MHC I | MHC II | MHC I & II | MHC I | MHC II | MHC I & II | MHC I | MHC II | MHC I & II | MHC I | MHC II | MHC I & II | MHC I | MHC II | MHC I & II |
| FLIDRINWI | 70.4% | 43.7% | 83.3% | 44.5% | 19.8% | 55.5% | 51.4% | 29.8% | 65.9% | 71.0% | 33.8% | 80.8% | 85.4% | 30.7% | 89.9% |
| FAYSHLERI | 74.9% | 40.2% | 85.0% | 50.6% | 31.8% | 66.3% | 60.9% | 40.5% | 76.7% | 77.5% | 35.5% | 85.5% | 89.2% | 19.5% | 91.3% |
| TVYHCSAVY | 61.3% | 40.1% | 76.8% | 54.5% | 18.4% | 62.9% | 63.7% | 45.0% | 80.0% | 50.4% | 43.5% | 72.0% | 50.2% | 20.9% | 60.6% |

**Table 15:**
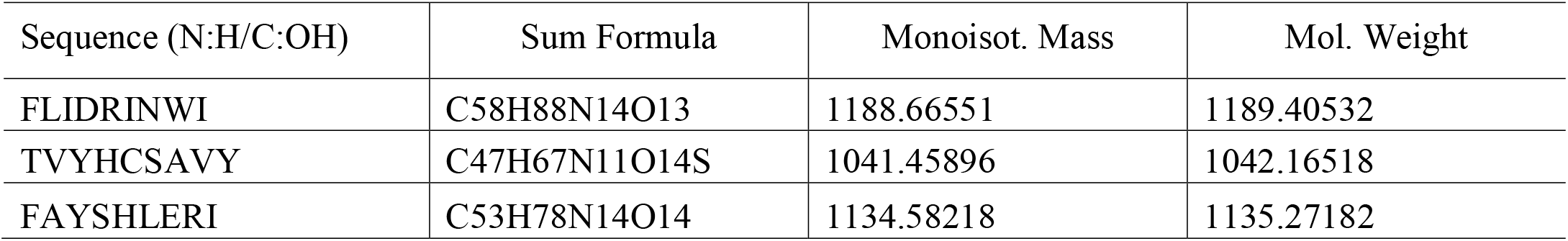
Monoisotopic Mass, Sum Formula and Molecular Weight of the three highly proposed peptides.

| Sequence (N:H/C:OH) | Sum Formula | Monoisot. Mass | Mol. Weight |
| --- | --- | --- | --- |
| FLIDRINWI | C58H88N14O13 | 1188.66551 | 1189.40532 |
| TVYHCSAVY | C47H67N11O14S | 1041.45896 | 1042.16518 |
| FAYSHLERI | C53H78N14O14 | 1134.58218 | 1135.27182 |

### 3.6. 3D Structure

**Figure 6:**
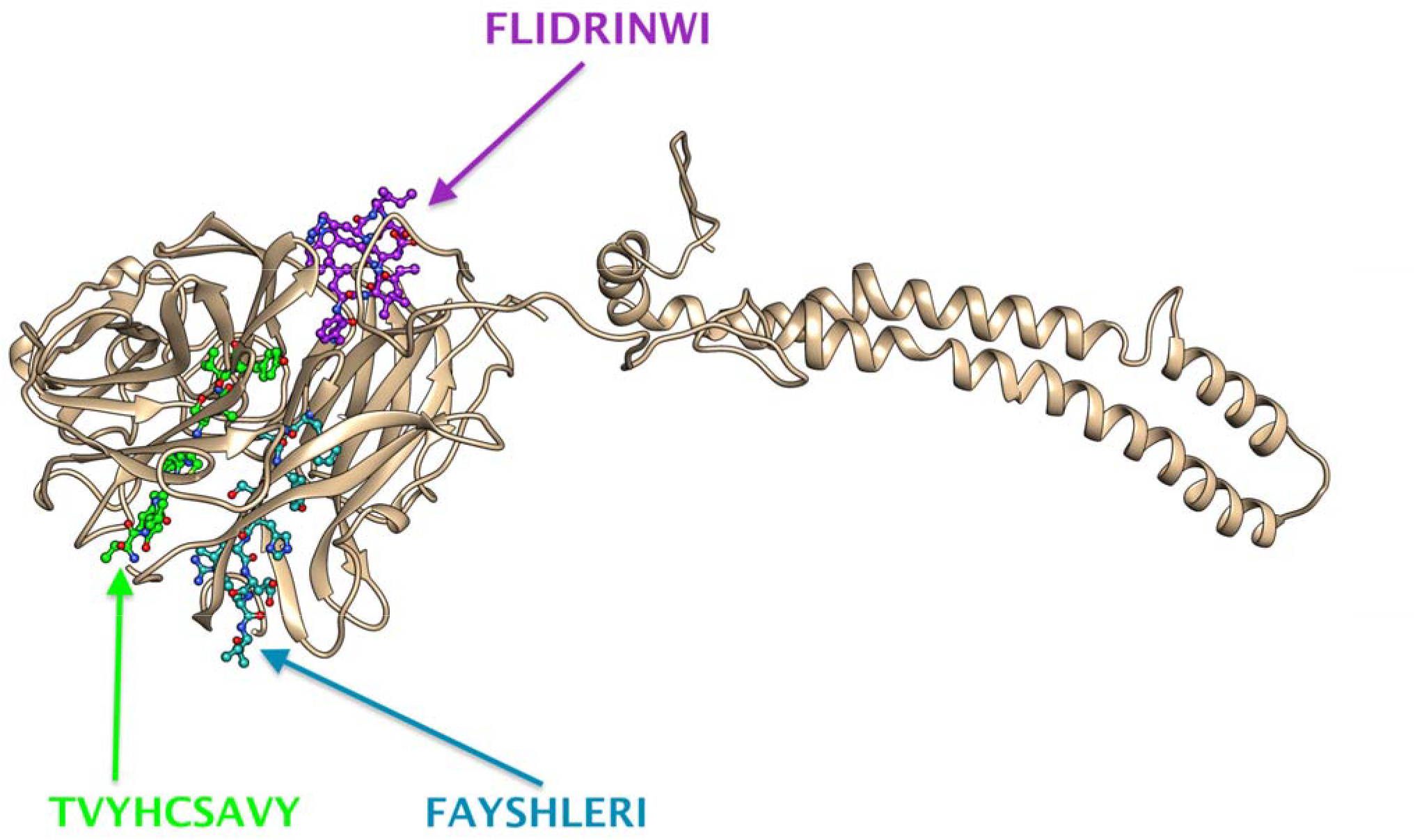
The four potential peptides bound to MHC I & MHC II visualized by Chimera X version 0.1.0.

### 3.7. Molecular Docking

**Figure 7:**
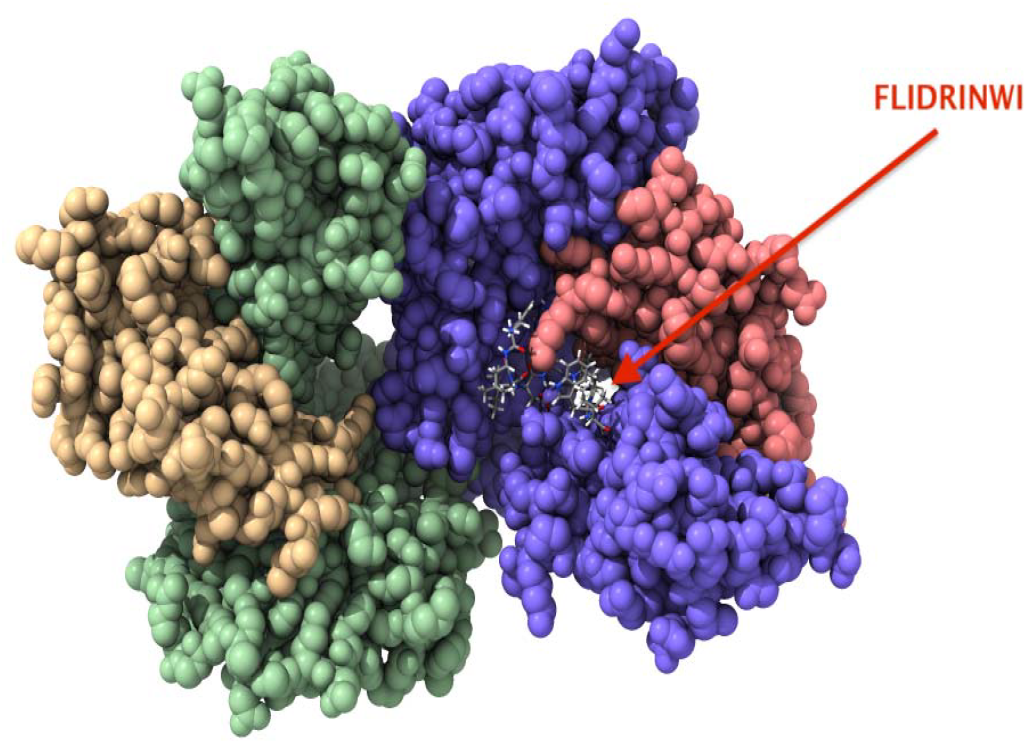
Molecular docking of FLIDRINWI peptide of Nipah virus docked in HLA-A0201 and visualized by UCSF Chimera X version 0.1.0.

**Figure 8:**
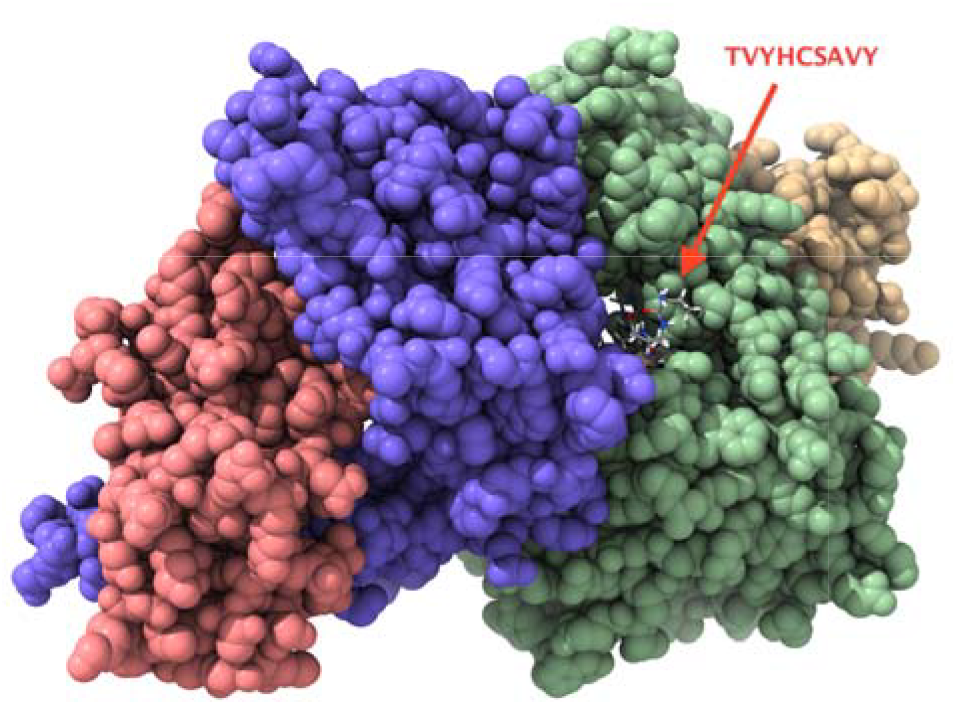
Molecular docking of TVYHCSAVY peptide of Nipah virus docked in HLA-A0201 and visualized by UCSF Chimera X version 0.1.0.

**Figure 9:**
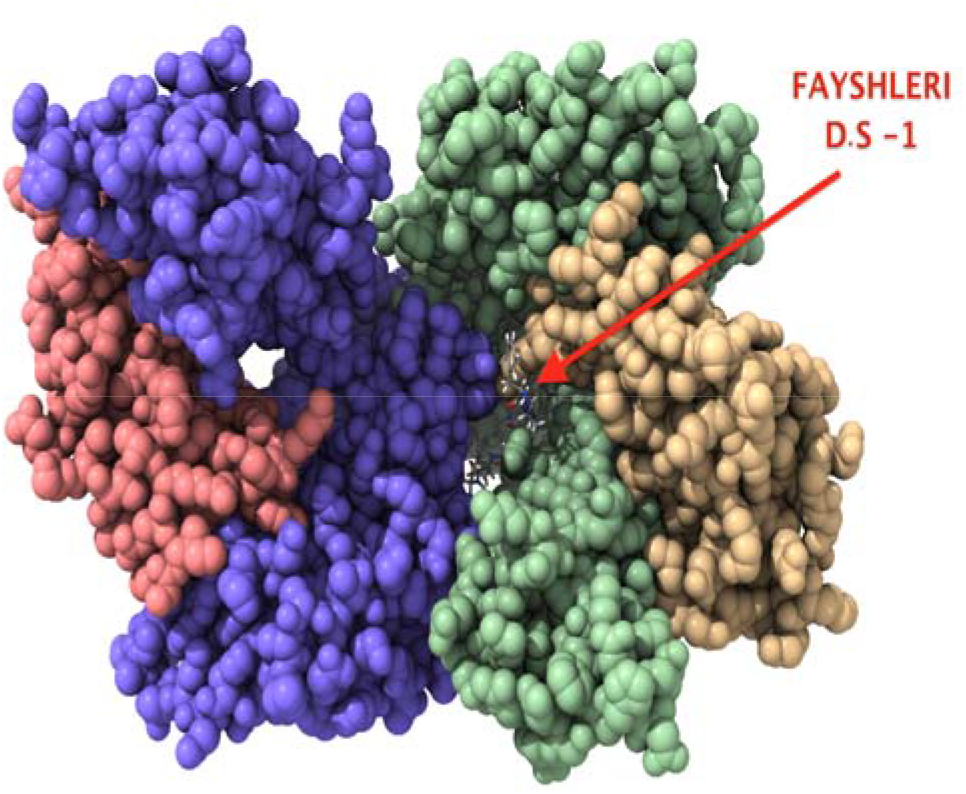
Molecular docking of FAYSHLERI peptide of Nipah virus docked in HLA-A0201 and visualized by UCSF Chimera X version 0.1.0. *D.S: Docking Side No.1.

**Figure 10:**
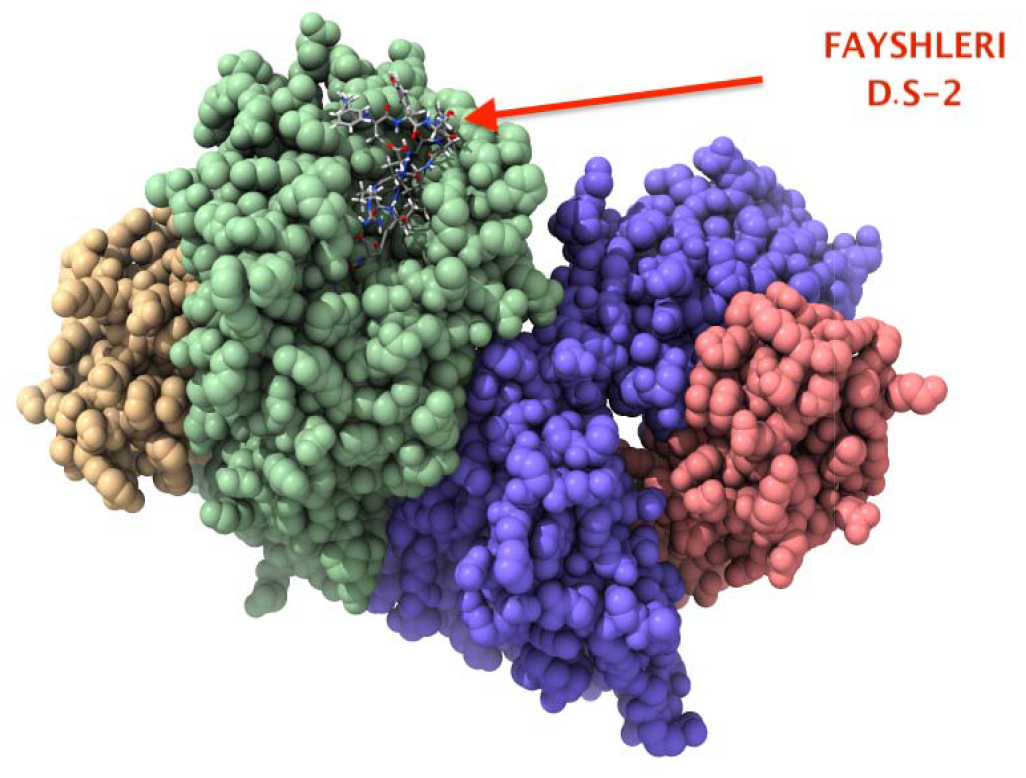
Molecular docking of FAYSHLERI peptide of Nipah virus docked in HLA-A0201 and visualized by UCSF Chimera X version 0.1.0. *D.S: Docking Side No.2.

## 4. Discussion

In our current study, potential peptides were suggested to design an epitope-based peptide vaccine against *Nipah virus glycoprotein G*. Literature was surveyed to define the antigenic part of the virus. Glycoprotein G that locates on the outer surface of the virus was selected to be our immunogenic target, those peptides were (FLIDRINWI, FAYSHLERI, TVYHCSAVY). FLIDRINWI was found to interact with 8 alleles in MHC I and 12 deferent alleles in MHC II with 83.3% MHC I & II World Population coverage. FAYSHLERI interacted with 11 alleles in MHC I and 13 alleles in MHC II while, the world population coverage of both MHC I & II were found to be 85.0%. Peptide TVYHCSAVY was found to interact with 11 alleles in MHC I and four alleles in MHC II while the MHC I & II world population coverage were found to be 76.8% (Table 2,3,14). Those three promising peptides showed a high susceptibility to be the first proposed vaccine against *Nipah virus glycoprotein G*.

We start by evaluating the binding affinity of MHC I alleles, by submitting the reference sequence of *Nipah virus glycoprotein G* to IEDB MHC I binding prediction tool based on the ANN align method with (IC_50_) ≤ 500 ^[37]^. 191 peptides were found to bind MHC I with different affinities. It is well known that a better immune response depends on the successful recognition of epitopes by HLA alleles with significant affinity. However, a peptide recognized by the highest number of HLA alleles has the best potential to induce a strong immune response. Therefore taking that fact, only three peptides were selected based on a higher number of alleles and 100% conservancy. The conserved peptide FLIDRINWI was found to interact with 8 alleles (HLA-A*02:01, HLA- A*02:03, HLA-A*02:06, HLA-A*68:02, HLA-C*03:03, HLA-C*06:02, HLA-C*07:01, HLA-C*12:03), while FSWDTMIKF interact with another 8 alleles (HLA-A*02:06, HLA-A*29:02, HLA-B*35:01, HLA-B*46:01, HLA-B*53:01, HLA-B*57:01, HLA-B*58:01, HLA-C*12:03) and TVYHCSAVY interact with 11 alleles (HLA-A*03:01, HLA-A*11:01, HLA-A*26:01, HLA- A*29:02, HLA-A*30:02, HLA-A*68:01, HLA-B*15:01, HLA-B*15:02, HLA-B*35:01, HLA-C*12:03, HLA-C*14:02), respectively. The reference sequence of *Nipah virus glycoprotein G* was analyzed again using the IEDB MHC II binding prediction tool based on NN-align with half-maximal inhibitory concentration (IC50) ≤ 1000; ^[38]^ analysis of binding to MHC II alleles resulted in prediction of 398 peptides from which FLIDRINWI, FAYSHLERI, FIEISDQRL and ILSAFNTVI were potentially proposed according to the high number of binding alleles. FLIDRINWI interact with 12 alleles, while FAYSHLERI interacts with 13 alleles, FIEISDQRL interacts with 17 alleles and ILSAFNTVI interact with 13 alleles. The population coverage results for total peptides binding to MHC1 alleles revealed 99.84% projected population coverage for the world, 98.55% in Southeast Asia, 98.40% in South Asia, 99.23% in North Africa and 99.36%in Sudan as presented on (Table 4-8), while the population coverage results for total peptides binding to MHC II alleles shown 56.84% projected population coverage for the world, 48.63%in Southeast Asia, 56.00%in South Asia, 62.37%in North Africa and 55.75% in Sudan (Table 9-13). The selected peptides were further subjected to both MHC I and MHC II based population coverage analysis in all World, Southeast Asia, South Asia, North Africa, and Sudan as shown on (Table 14). Among the five primarily selected peptide, the obtained results showed very strong potential in proposing peptide FLIDRINWI as vaccine candidate than others by considering its overall peptide conservancy, population coverage and by the affinity for the highest number of HLA alleles.

The peptide FLIDRINWI demonstrate a special population coverage results for both MHC I and MHC II alleles with total binding to 12 alleles in MHC II and 8 alleles in MHC I, This finding demonstrate a very strong potential to formulate an epitopes-based peptide vaccine against *Nipah virus glycoprotein G* and make the peptide FLIDRINWI highly proposed. Furthermore, an *in-silico* molecular docking was done to explore the binding affinity between the aforementioned peptides and the target HLA-A0201; which has been selected for docking pertained with its involvement in several immunological and pathological diseases ^[46,47,48]^. Although a numerous studies have demonstrated an association between HLA alleles and disease susceptibility, defining protective HLA allelic associations potentially allows the identification of pathogen epitopes that are restricted by the specific HLA alleles. These epitopes may then be incorporated into vaccine design in the expectation that the natural resistance can be replicated by immunization ^[47,48]^. Calculation of the root means square deviation (RMSD) between co-ordinates of the atoms and formation of clusters based on RMSD values computed the resemblance of the docked structures. The best docking result is considered to be the conformation with the lowest binding energy. The least energy was predicted for the peptide FLIDRINWI (−6.95 Kcal/mol) and the 3D structure of the allele and peptide shown in (figure 7). According to these interesting outcomes, formulating a vaccine using FLIDRINWI peptide is highly promising that is encouraging to be the first proposed epitope-based peptide vaccine against *Nipah virus*. However, novel vaccines approach as epitope-based peptide vaccines can possibly conquer these obstructions for these immunizations can make increasingly successful, specific and long lasting impregnable reaction and without all the adverse effects of the classical vaccines. Moreover, based on our previous studies peptide-based vaccines have been effectively proposed by utilizing in silico approach against *Madurella mycetomatis*, Mokola Rabies Virus, Lagos Rabies Virus and others ^[49,50,51]^. Such investigations in regards to those viruses built up immunoinformatics considered as a computational analysis. In this study the same techniques were used, to design the Epitope-Based Peptide Vaccine against *Nipah virus glycoprotein G* as a target immunogenic part of the virus to stimulate a protective immune response.

One of the limitations of this study was the absent of peptides with a strong binding affinity to B- cell The sequence of *Nipah virus glycoprotein G* was subjected to Bepipred linear epitope prediction, Emini surface accessibility, and Kolaskar and Tongaonkar antigenicity methods in IEDB.

## 5. Conclusions

To the best of our knowledge, this study is considered to be the first to propose epitope-based peptide vaccine against glycoprotein G of Nipah virus, which is expected to be highly antigenic with a minimum allergic effect. Furthermore, this study proposes promising peptide FLIDRINWI with a very strong binding affinity to MHC1 and MHC11 alleles. This peptide shows an exceptional population coverage results for both MHC1 and MHC11 alleles.

In-vivo and in-vitro assessment for the most promising peptides namely; (FLIDRINWI), (TVYHCSAVY) and (FAYSHLERI) are recommended to be explored and find out more on their ability to be developed into vaccines against *Nipah virus glycoprotein G*.

